## Supplemental material for "Using colony size to measure fitness in *Saccharomyces cerevisiae*"

| Strain | Description | Source |
|---|---|---|
| COPR_A EH0; YJF4668 | Evolved from ancestors d1E1 and d2E1 in CM. | [34] |
| COPR_A EH0_80; YJF4669 | Evolved from ancestors d1E9 and d2E9 in CM and CM + 6.4 μM $CuSO_4$ on alternating days | |
| COPR_A EH80; YJF4670 | Evolved from ancestors d1H5 and d2H5 in CM + 6.4 μM $CuSO_4$ | |
| SaltA EH0; YJF4671 | Evolved from ancestors d1E1 and d2E1 in CM | |
| SaltA EH0_80; YJF4672 | Evolved from ancestors d1E9 and d2E9 in CM and CM + 274 mM NaCl on alternating days | |
| SaltA EH80; YJF4673 | Evolved from ancestors d1H5 and d2H5 in CM + 274 mM NaCl | |
| Ancestor d1E1; YJF4674 | Derived from mating YJF153 (MATa, HO::dsdAMX4 with barcoded kanMX deletion casettes from the MoBY plasmid collection) and YJF154 (MATalpha, HO::dsdAMX4). Both parents are derivatives of an oak tree strain, YPS163. | |
| Ancestor d2E1; YJF4675 | | |
| Ancestor d1E9; YJF4676 | | |
| Ancestor d2E9; YJF4677 | | |
| Ancestor d1H5; YJF4678 | | |
| Ancestor d2H5; YJF4679 | | |
| YJF4604 | YJF1389 (MATa, HO::YFP-NAT, ura3-140) mated to YJF154 (MATalpha, HO:dsdAMX4). Both parents are derivatives of YPS163. | This study, [35] |

**S1 Table. Strains used in this study.** Evolved strains are indicated by the name of their stress (COPR_A for copper and SaltA for sodium), and the percent lethal limit in which they were raised (*e.g.* EH0_80 indicates an evolutionary history of environment fluctuation between 0 and 80 percent lethal stress).

| Assay | Stress | Stress Concentration | Sample Number | RMSE | $R^2$ | Smallest Detected Fitness Difference | Minimum Detectable Fitness Difference |
|---|---|---|---|---|---|---|---|
| Colony Size | $CuSO_4$ | 0 μM | ~32 | 0.0085 | 0.629 | 0.0026 | 0.0043 |
| | | 10 μM | | 0.0086 | 0.485 | 0.0030 | 0.0044 |
| | | 20 μM | | 0.0281 | 0.909 | 0.0130 | 0.0144 |
| | | 30 μM | | 0.1128 | 0.859 | 0.0451 | 0.0577 |
| | | 40 μM | | 0.2138 | 0.869 | 1.1021 | 0.1093 |
| | NaCl | 0 mM | | 0.0107 | 0.518 | 0.0041 | 0.0055 |
| | | 10 mM | | 0.0108 | 0.430 | 0.0024 | 0.0055 |
| | | 50 mM | | 0.0076 | 0.678 | 0.0025 | 0.0039 |
| | | 100 mM | | 0.0087 | 0.847 | 0.0037 | 0.0044 |
| | | 150 mM | | 0.0153 | 0.685 | 0.0079 | 0.0078 |
| | | 200 mM | | 0.0121 | 0.743 | 0.0050 | 0.0062 |
| | | 300 mM | | 0.0176 | 0.724 | 0.0361 | 0.0090 |
| | | 400 mM | | 0.0294 | 0.718 | 0.0147 | 0.0150 |
| | | 500 mM | | 0.0171 | 0.836 | 0.0119 | 0.0088 |
| | | 600 mM | | 0.0190 | 0.874 | 0.0182 | 0.0097 |
| | | 800 mM | | 0.0404 | 0.565 | 0.0145 | 0.0207 |
| | | 1000 mM | | 0.0239 | 0.498 | 0.0131 | 0.0122 |
| | | 1200 mM | | 0.0461 | 0.244 | 0.0311 | 0.0236 |
| Comp-etitive Fitness | CM | 0 μM | 7 | 0.0107 | 0.903 | 0.0221 | 0.0136 |
| | $CuSO_4$ | 5 μM | | 0.0168 | 0.779 | 0.0050 | 0.0214 |
| | NaCl | 103 mM | | 0.0145 | 0.724 | 0.0148 | 0.0185 |

**S2 Table. Summary of measurement properties from the competitive fitness and colony size assays.** The root mean square error (RMSE) and variance in fitness explained by strain ($R^2$) are derived from the fitness difference ~ evolved strain relationship at each stress concentration using the data shown in Figs 2 and 3. Minimum detectable fitness differences is from power analysis at 0.80 power and 0.95 significance.

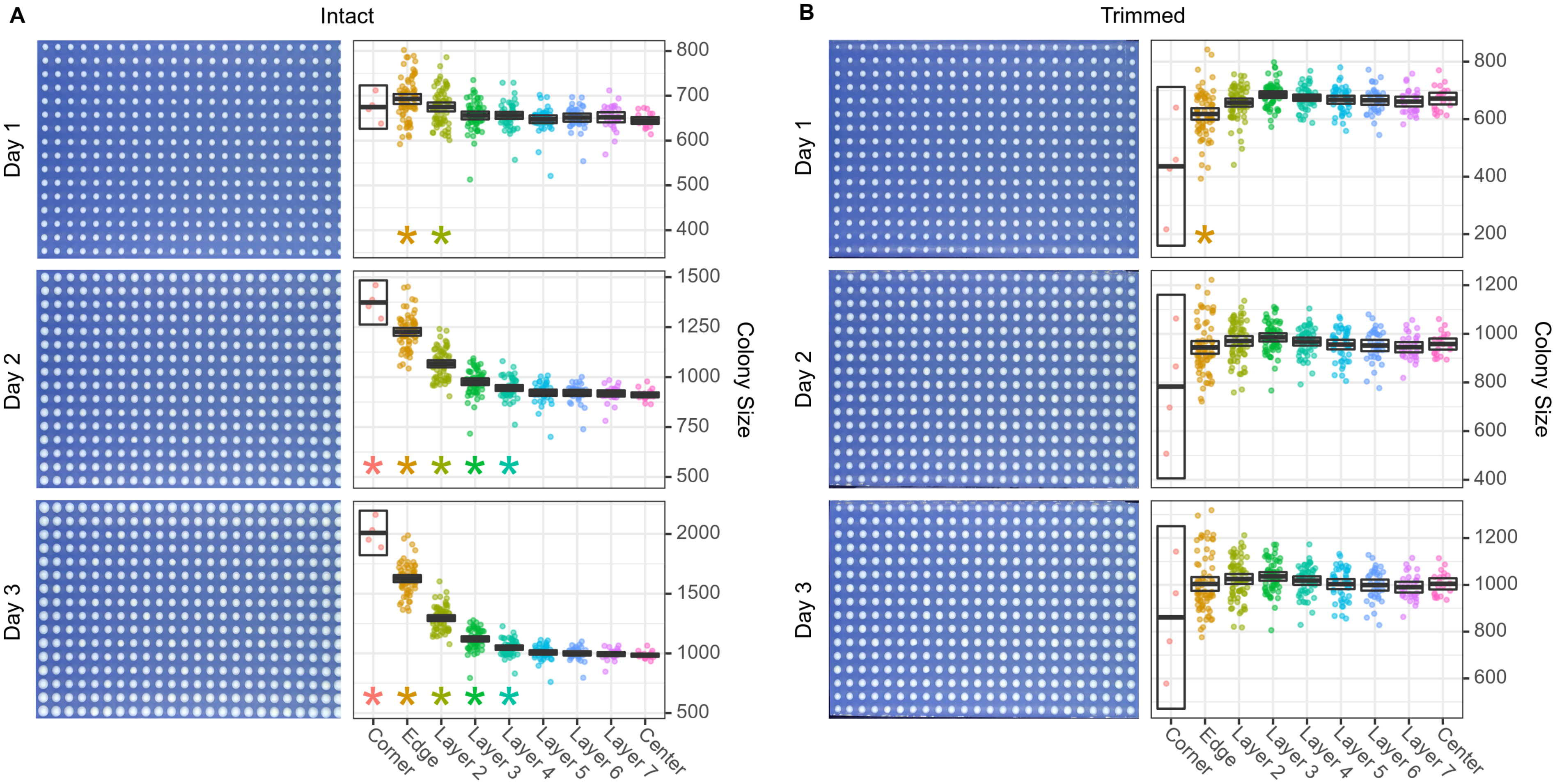


**S1 Figure. Edge effects over time.** Colony photographs and size distributions by layer for days 1-3 associated with Fig 1B-E. Evolved and ancestor strains were arrayed in a randomized pattern on CM agar plates either left intact or trimmed immediately after printing (A and B, respectively). Boxes indicate the 95% confidence interval bisected by the mean. Stars indicate layers that are significantly different from the center (FDR-corrected 2 sample t-test).

**A**

| | Ancestor d1H5 | Ancestor d2H5 | COPR_A EH80 | SaltA EH80 |
|---|---|---|---|---|
| Day 1 | 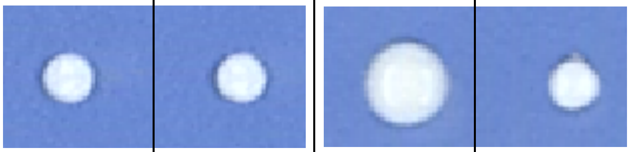 | | | |
| Size | 237 | 231 | 687 | 247 |
| Cells | 4.667E+06 | 4.495E+06 | 2.217E+07 | 4.958E+06 |
| Average Ancestor Generations | | | 2.9826 | |
| log(Cells)~Generations | 0.699 | 0.687 | 1.222 | 0.720 |
| Fitness Difference | 0.0063 | -0.0063 | 0.5287 | 0.0266 |
| Day 2 | 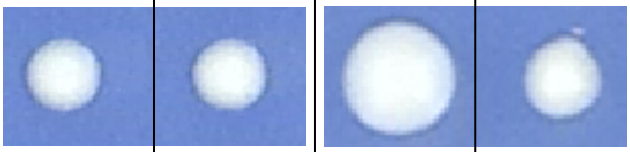 | | | |
| Size | 500 | 496 | 1292 | 595 |
| Cells | 1.392E+07 | 1.376E+07 | 5.589E+07 | 1.796E+07 |
| Average Ancestor Generations | | | 4.578 | |
| log(Cells)~Generations | 0.695 | 0.691 | 1.027 | 0.746 |
| Fitness Difference | 0.0019 | -0.0019 | 0.3334 | 0.0530 |
| Day 3 | 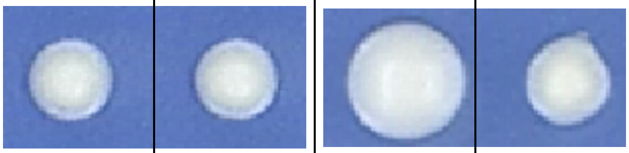 | | | |
| Size | 692 | 683 | 1524 | 780 |
| Cells | 2.240E+07 | 2.198E+07 | 7.117E+07 | 2.670E+07 |
| Average Ancestor Generations | | | 5.259 | |
| log(Cells)~Generations | 0.695 | 0.692 | 0.934 | 0.737 |
| Fitness Difference | 0.0014 | -0.0014 | 0.2411 | 0.0438 |
| Day 4 | 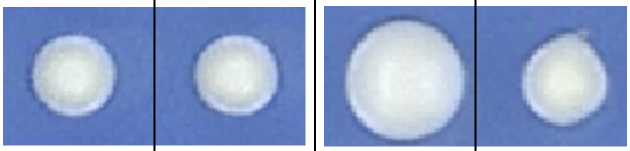 | | | |
| Size | 714 | 705 | 1567 | 809 |
| Cells | 2.346E+07 | 2.302E+07 | 7.413E+07 | 2.816E+07 |
| Average Ancestor Generations | | | 5.3255 | |
| log(Cells)~Generations | 0.694 | 0.692 | 0.902 | 0.734 |
| Fitness Difference | 0.0012 | -0.0012 | 0.2084 | 0.0411 |

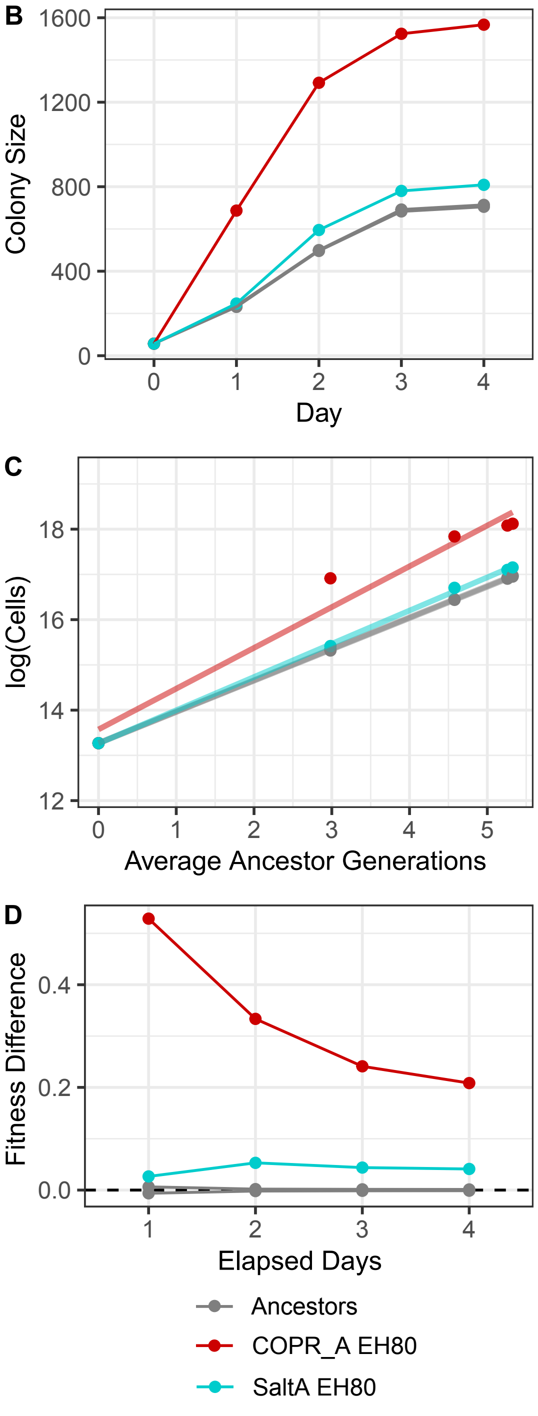


**S2 Figure. Estimating fitness differences from colony size.** Examples of colony size, estimated number of cells, generations and fitness differences over four days with corresponding photographs (A). Colony size areas (pixels) were recorded for four consecutive days after pinning and the size on day 0 was set to 57 (B). Colony size was converted to cell number using the experimentally derived log-log relationship (Fig 1A) and plotted as a function of ancestor generations (C). Fitness differences were determined from the difference in regression slopes (from panel C) between each evolved strain and the average of its ancestor pair as a function of elapsed days (D).

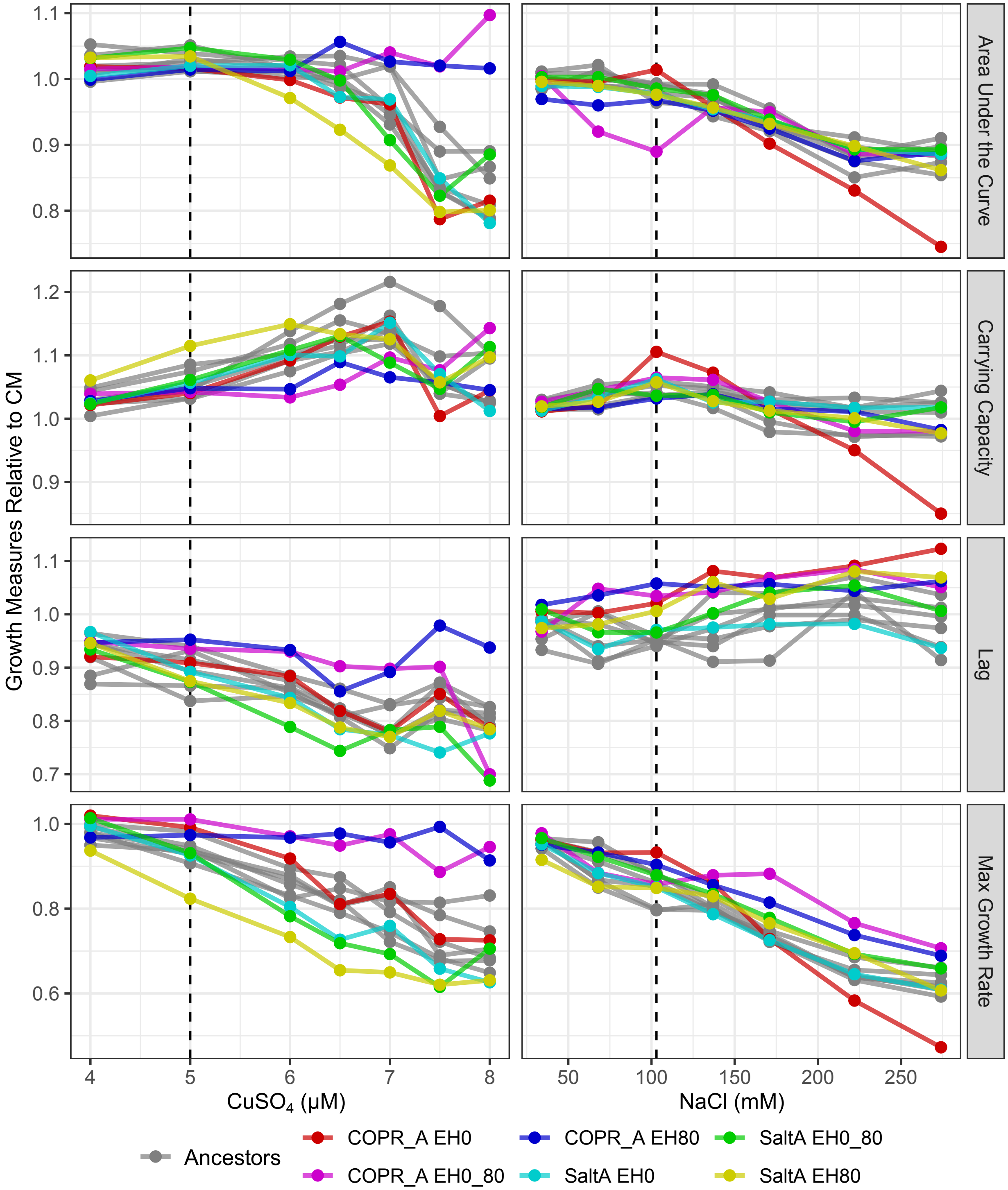


**S3 Figure. Liquid growth curve characteristics of evolved and ancestor strains.** Pure cultures were used to determine the single stress concentrations for competitive growth assays. The area under the curve, carrying capacity, time lag and maximum growth rate for different concentrations of copper and sodium were determined from growth curves of each strain (n = 1). Growth measures are shown relative to growth in CM without added copper or sodium. Dotted lines indicate the concentration chosen for competitive fitness assays.

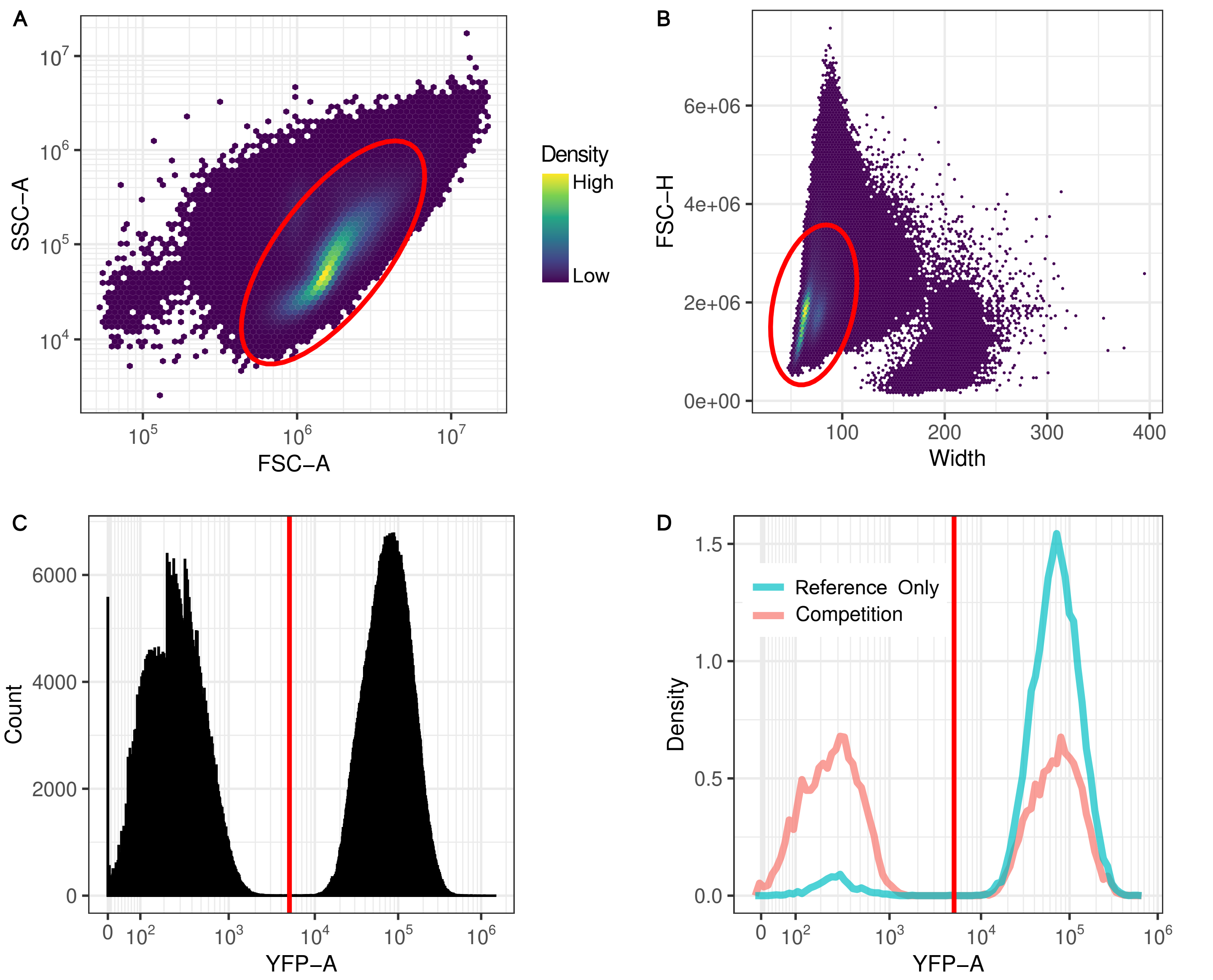


**S4 Figure. Gating scheme used to measure the proportion of YFP cells for the competitive fitness assays.** Gating parameters were determined from the pooled events of all runs pertaining to a single day and stress. The first gate excluded events outside the 99% ellipse from log10 SSC-A (side scatter area) vs. log10 FSC-A (forward scatter area)(A). This was followed by event exclusion from a 95% ellipse on a FSC-H (forward scatter height) vs. width (B). The YFP gate was manually set at the midpoint between the two largest populations on a log10 YFP-A histogram and kept at a constant value throughout all experiments (C). Example YFP-A histograms after gating showing the competition between an ancestor strain and the YFP-expressing reference strain, and a reference-only control consisting of YJF4606 alone (D). Note the small proportion of the events exhibited fluorescent signalling below the YFP gate in these reference only controls (1-19%) which are accounted for as the false negative fraction (Equation 1).

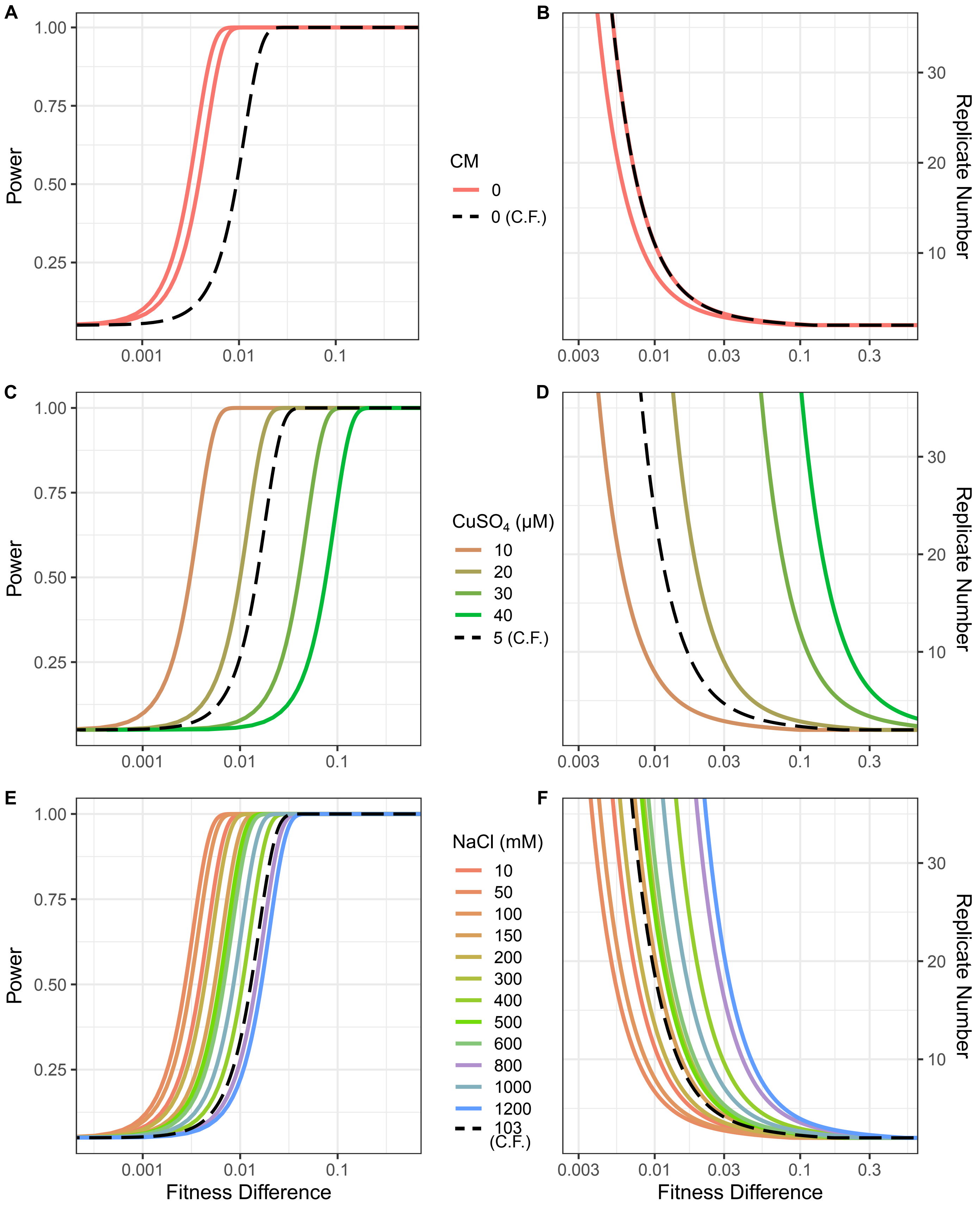


**S5 Figure. Power analyses of competitive fitness and colony size assays.** Power and number of replicates needed for competitive fitness (dashed lines, labeled "C.F.") and colony size assays (solid lines) for CM only (A and B) and various concentrations of copper (C and D) and sodium (E and F). Power as a function of fitness difference was derived from each assay's measured performance (A, C and E). The number of replicates needed to achieve 0.8 power at 0.95 significance is plotted as a function of fitness difference (B, D and F).

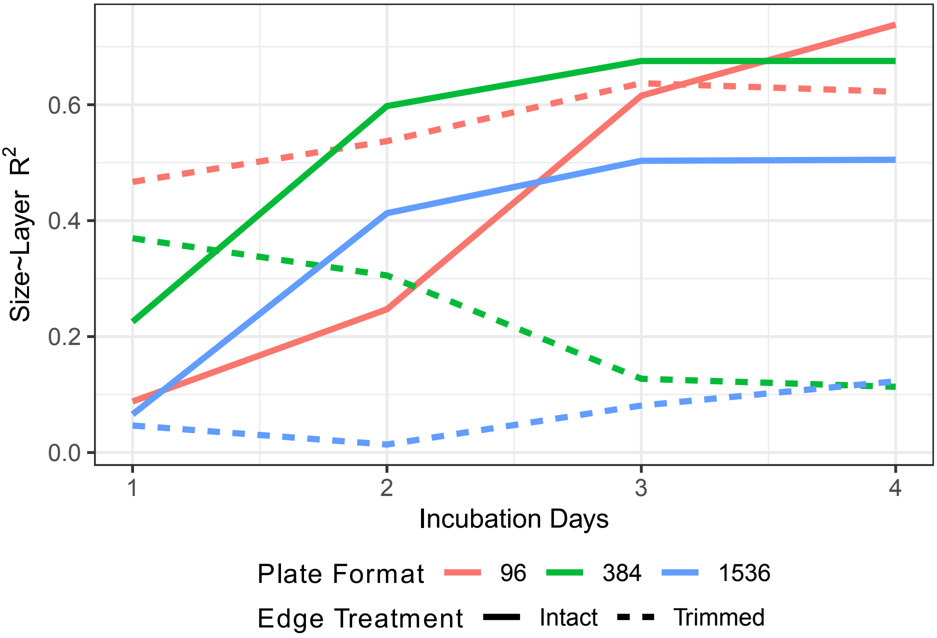


**S6 Figure. The proportion of variance explained by layer as a function of incubation time.** Variance explained ($R^2$) is shown for intact and trimmed plates arrayed on CM plates in 96, 384 or 1536 density formats. All colonies are of ancestor strain YJF4679 and corner colonies were excluded.

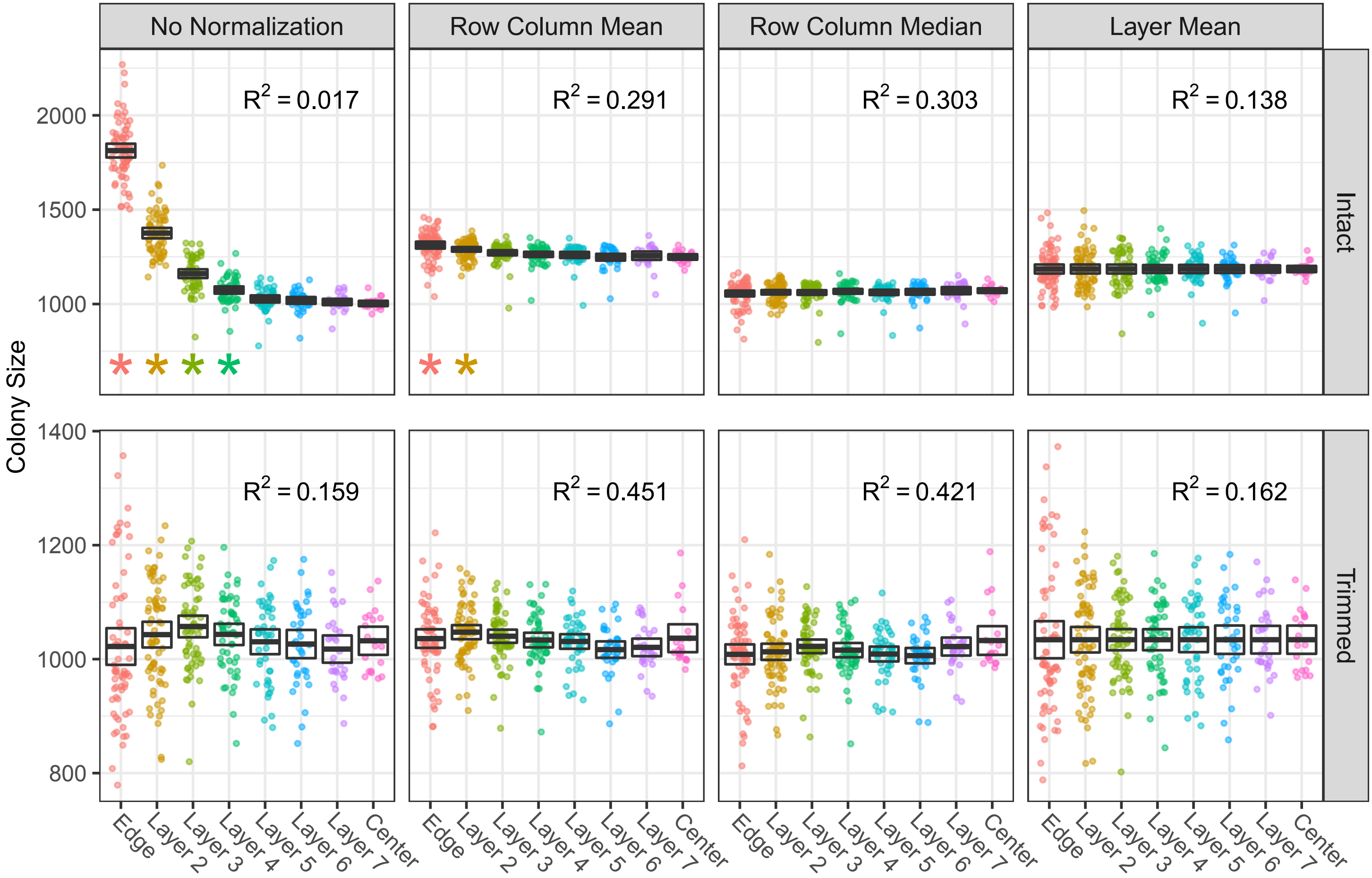


**S7 Figure. Colony sizes from various normalization methods.** Colonies are after 4 days growth from trimmed and intact CM plates with no normalization and normalization by row and column mean, row and column median, or layer. Numbers in the upper right indicate the size variance explained by strain (colony size ~ strain). Boxes show the 95% confidence interval bisected by the mean with stars indicating layers that significantly differed from the center layer (FDR-corrected 2 sample t-test).

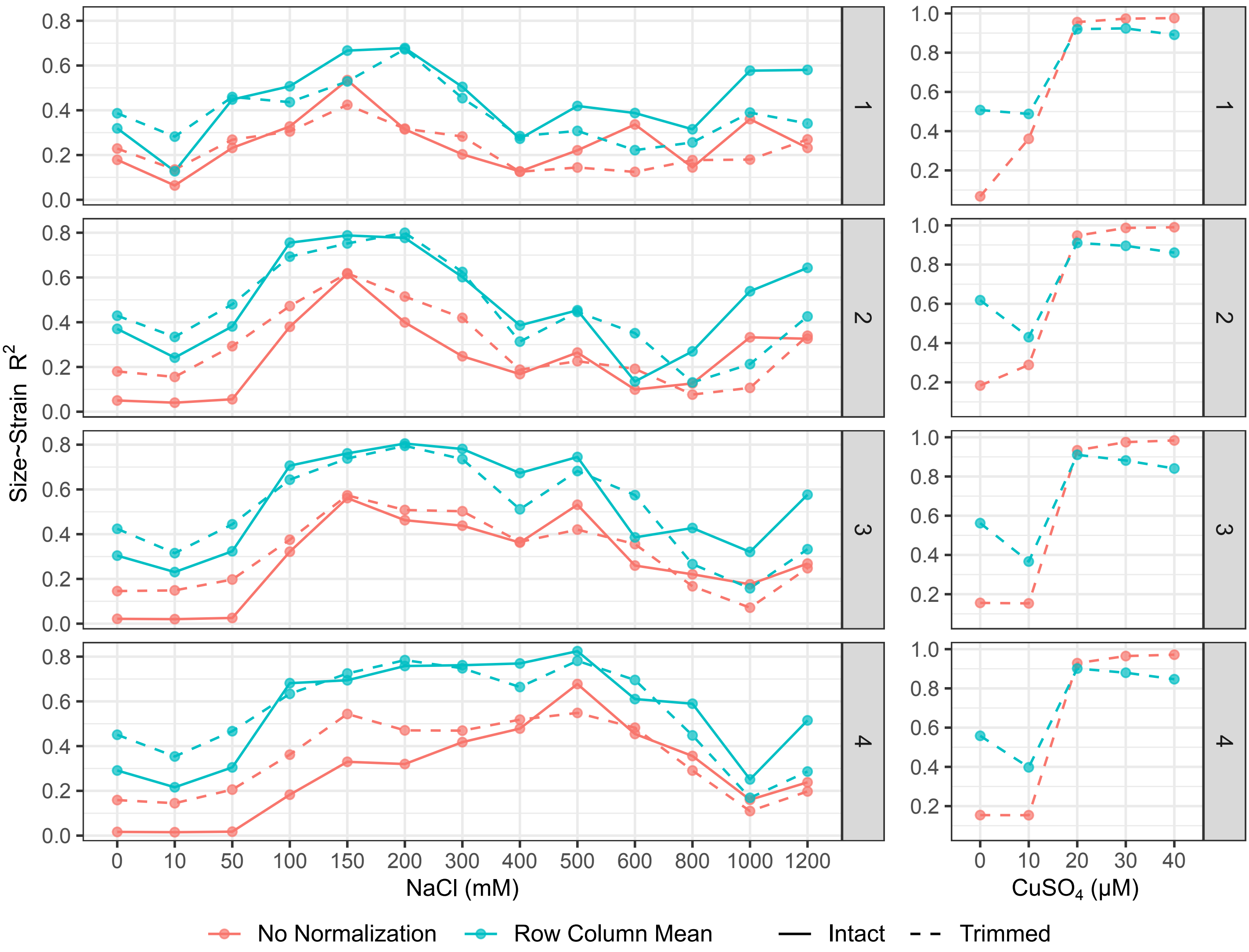


**S8 Figure. Colony size variance explained by strain after normalization.** Variance explained by strain ($R^2$) before and after row and column mean normalization for either intact or trimmed plates containing various concentrations of sodium (left) or trimmed plates with copper (right).

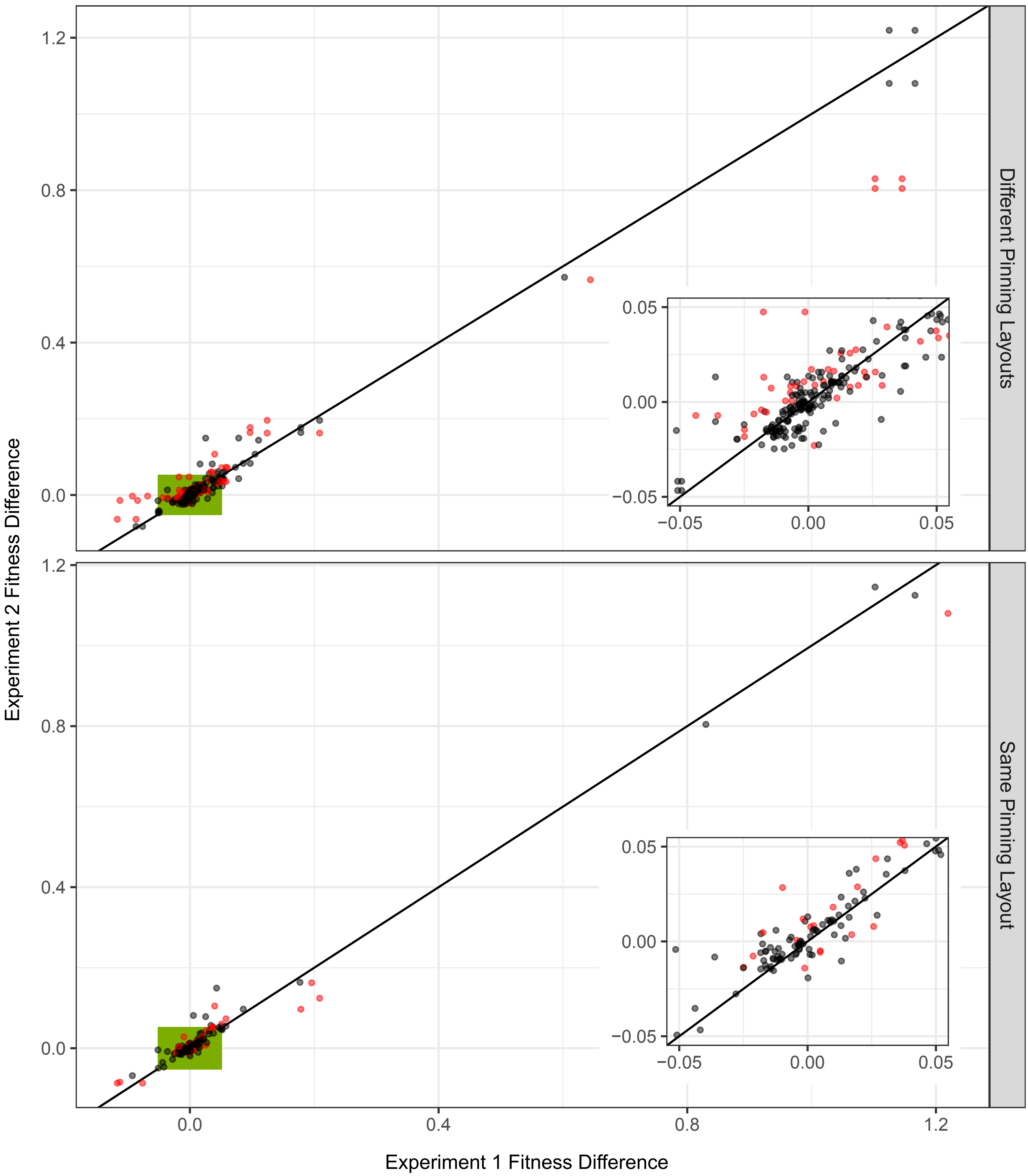


**S9 Figure. Fitness differences estimated from independent experiments with either same or different pinning layouts.** Points in black indicate that independent experiments resulted in overlapping 95% confidence intervals whereas points in red did not. Green boxes indicate the locations of the insets shown in each plot, the y=x line is shown in black.

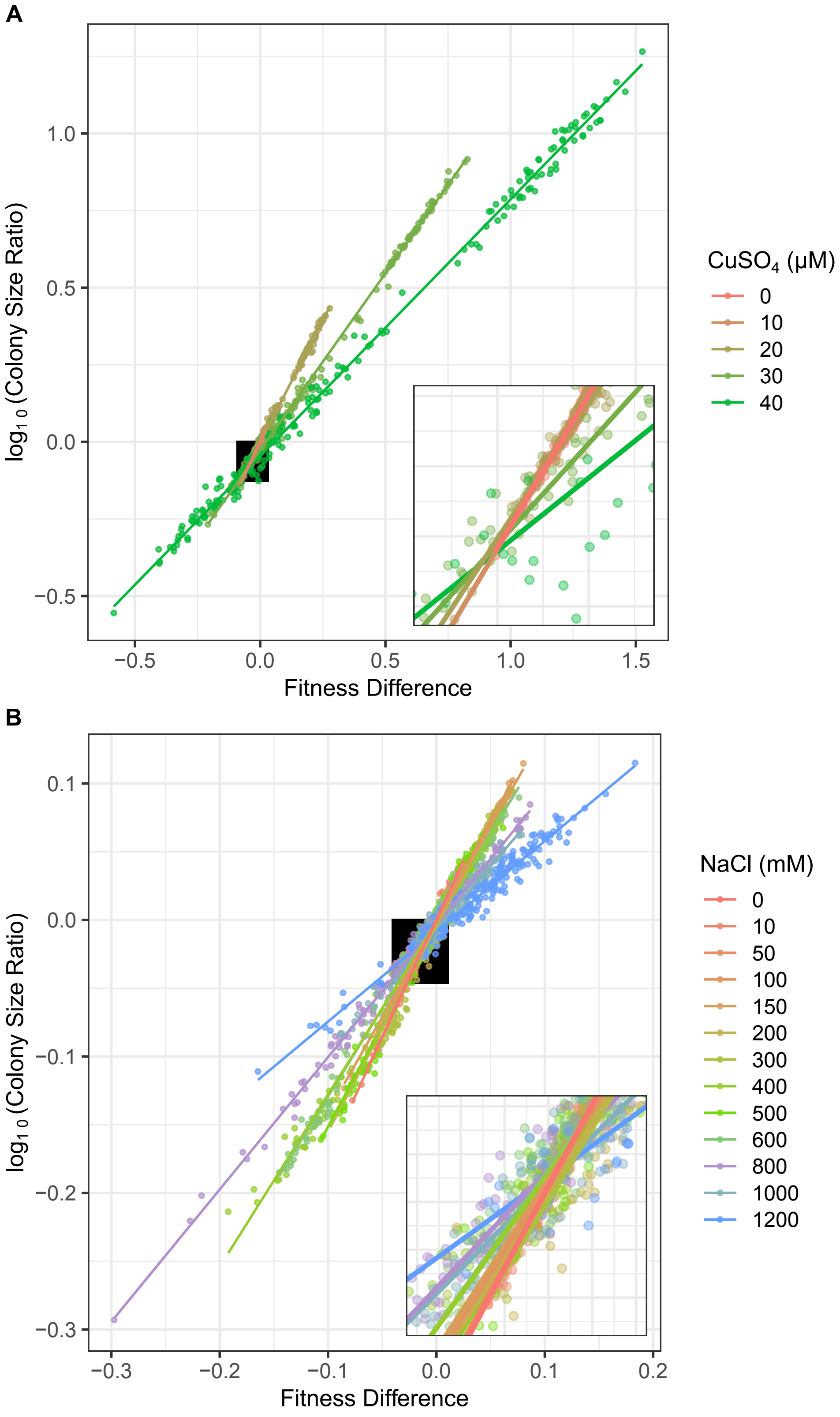


**S10 Figure. Comparison of fitness differences and colony size ratios.** The log of colony size ratios from Fig 3 were calculated by dividing the colony size of each evolved replicate by the average size of its respective ancestors on copper (A) and sodium (B) plates. The location of the inset is indicated by the black box.
